# Departures from Mendelian inheritance of *EPSPS* gene copies in a herbicide-resistant weed

**DOI:** 10.1101/2025.05.12.653577

**Authors:** Lisa H. Han, Kathryn F. Dungey, S.B. Yakimowski

## Abstract

Mendelian transmission requires a stable genetic structure, but unstable elements contribute to dynamic genomes. Here, we report on non-Mendelian inheritance and intra-individual instability of extrachromosomal gene copies, which are associated with rapid evolution of herbicide resistance in the dioecious, agricultural weed *Amaranthus palmeri*. Copy number variation for the glyphosate resistance gene *EPSPS* was quantified using digital droplet PCR within individuals, and for parents and offspring of controlled crosses with copy number ranging from 1 to ∼160. Parent-offspring regressions of copy number indicates non-Mendelian inheritance. Heritability is higher (*h*_G_^2^=1.60) below ∼33 copies, than above (*h*_G_^2^=0.22) because parents with fewer gene copies yield offspring with similar or increased mean copy number, whereas offspring of high copy number parents have fewer copies than parents. Intra-individual variance in copy number increased with mean individual copy number, likely contributing to variation observed in offspring copy number, but not sufficient to explain the most extreme transgressive copy number variants, nor the loss of high copy number. In short, copy number is less stable and less heritable in parents with more *EPSPS* copies, with offspring losing *EPSPS* copies. This suggests that genome instability imposes a constraint on the evolution of herbicide resistance through increased *EPSPS* copy number.

## 1. Introduction

Modern biology is largely built around a Mendelian model of inheritance, as described by the Modern Synthesis [1,2]. But, this view is complicated by structural genomic re-arrangements among individuals [3], and plastic genomic re-organization in response ‘to stress or unusual challenge’ as investigated by McClintock (e.g. transposable elements) in the early decades of genetics [4]. Variation in gene copy number represents polymorphism in genome structuring, and is observed in humans and many other species [3,5,6], including model plant *Arabidopsis thaliana* for which copy number varies for ∼one-third of genes [7]. Copy number variation can affect key phenotypes including fitness. For example, tandem repeats of multiple genes in soybean confers nematode resistance [8], and copy number variation of two genes is associated with flowering time variation in wheat (*Triticum aestivum* L.) [9–11]. More recent studies have demonstrated the occurrence of gene copies residing on ‘extra-chromosomal DNA’ [12,13], distinct from the core genome. Yet, in the current era of high-throughput sequencing and bioinformatics, the methods required to resolve structural genomic variation are more complex [14–17] and thus understanding of the dynamic nature of genomes has lagged behind study of relatively stable genome sequence variants [18]. In turn, our understanding of the evolutionary dynamics [19] of multi-copy gene traits [20–22] remains limited.

Rapid evolution of herbicide resistance is a threat to food security and the sustainability of agricultural practices [9,10]. At least 60 plant species worldwide have reported resistance to the ubiquitous herbicide glyphosate [23] since its commercialization in 1974 [24,25]. A variety of genetic mechanisms of resistance have been detected; for example, point mutations and increased copy number [12,26] of the *EPSPS* (5-enolpyruvylshikimate-3-phosphate synthase) gene have resulted in the functional maintenance of the shikimate pathway under glyphosate stress [27–29]. The North American weed *Amaranthus palmeri* (Palmer’s Amaranth) has evolved extrachromosomal positioning of highly duplicated genes. This extrachromosomal DNA (ecDNA) is often circular (eccDNA) and putatively tethers to chromosomes, replicating and segregating during mitosis and meiosis [30,31]. The discovery of ecDNA gene copies is relatively recent in plants [13,30,32], but well known in bacteria (i.e. plasmids) since plasmids were described in the 1960s (e.g. *penicillinase* plasmids in *staphylococcal* strains -Staph. Aureus) [33–35]. The potential instability of extrachromosomal gene copy number raises fundamental questions about the genomic basis of inheritance and the influence of environmentally-induced genome plasticity on rapid evolution.

Genome instability violates assumptions of Mendelian inheritance, resulting in deviations from expectations of Mendel’s laws of segregation and assortment. Heritability is a fundamental concept in genetics and evolutionary change [36] that extends Mendel’s laws to polygenic traits [37,38] as described in the Modern Synthesis of Evolutionary Biology [1]. Heritability quantifies the genetic and environmental contributions to the expression of a quantitative trait. This can apply to an inherited trait, including genome structure itself. In particular, estimation of heritability based on the resemblance between parents and offspring can also be applied to copy number of an individual gene [39,40], facilitating a more direct estimate [36] of transmission of a specific gene, and any potential departures from Mendelian inheritance.

Extrachromosomal DNA is an example of genetic material defined by unique characteristics relative to chromosomal DNA, thus giving rise to questions about its pattern of variation through reproduction. Extrachromosomal circular DNA (‘eccDNA’) was first described in 1965 [41] and is a general term for double-stranded DNA that resides external to the chromosomes, varying in size from smaller (0.2 - 100kb) to larger structures (several hundred kb to several Mb). It has been identified in both healthy cells [42,43] and cancerous tumour tissue [44], and varies in content (e.g. repetitive and transcribed regions) [45,46]. Extrachromosomal DNA was detected in plants 20 years later, but recent genomic and bioinformatic methods have revealed ec(c)DNA in several model plants (e.g. *Arabidopsis* and *Poplar*) and agriculturally-important species (e.g. rice, potato, carrot). The agricultural weed *Amaranthus palmeri* has the largest extrachromosomal structure (∼400kb replicon) detected in plants to date [47]. In contrast to many small, non-coding eccDNAs, the *A. palmeri* replicon carries a copy of the glyphosate resistance gene (*EPSPS*) and >50 other genes, most of which are transcribed [31]. This well-characterized structure of *A. palmeri* eccDNA presents an opportunity to investigate inheritance and stability of gene copy number for a transcribed, extrachromosomal resistance gene.

Here, we study inheritance and stability of resistance gene copy number transmission in *A. palmeri* (Palmer Amaranth), a crop weed with reported resistance to ten herbicidal modes of action [23,48], including glyphosate. The glyphosate resistance phenotype in *A. palmeri* is strongly associated with *EPSPS* copy number variation [49], with copy number ranging from one to over 160 per genome [30,31]. Experimental crosses between single/low copy x high copy number individuals established that eccDNA can be transmitted from parent to offspring. However, fiber-FISH analysis of copy number variation revealed variation in the number and position of eccDNAs following mitosis [30]. This cellular variation in eccDNA raises the question of whether parental copy number is heritable. To investigate the stability of a resistance gene copy number polymorphism with extrachromosomal positioning we estimated heritability with 39 crosses (n=1041 offspring total) and intra-individual variation for 117 plants (n=749 total estimates) of *EPSPS* copy number spanning the dynamic range of *EPSPS* copy number observed in the multi-resistant agricultural weed *Amaranthus palmeri*. Drawing from quantitative genetics, we address the following questions: (1) Does midparent copy number predict mean offspring copy number? If so, is the heritability relationship linear and 1:1, as predicted by the Mendelian expectation for the segregation and assortment of copy number variation? (Fig. 1A). (2) Are deviations of offspring from mean parental copy number randomly distributed around a constant mean (∼0), and independent of parental midpoint copy number, as expected for measurement error (Fig. 1C)? Or, is there evidence for copy number instability in the form of greater deviations between offspring and parents as copy number increases? (3) How does intra-individual variation in *EPSPS* copy number compare to inter-individual and parent-offspring copy number variation? In particular, does genome instability affect copy number throughout growth and development, yielding directional loss or gain of copies?

**Figure 1.**
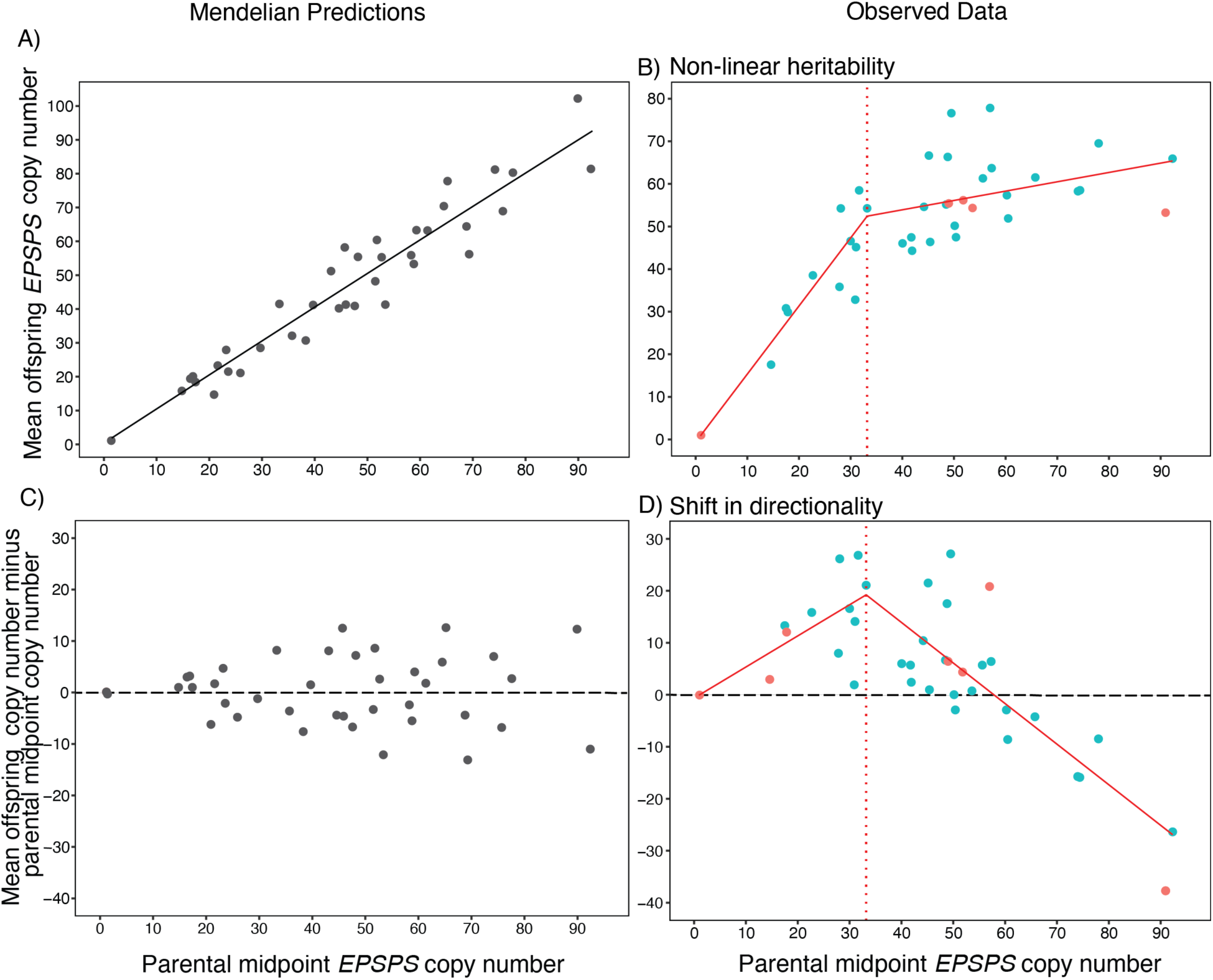
A) Predicted linear and ∼1:1 relation between parental midpoint EPSPS copy number and mean offspring copy number under Mendelian inheritance. B) Observed non-linear relation between parental midpoint *EPSPS* copy number and mean offspring EPSPS copy number in *Amaranthus palmeri* (n= 39 crosses). C) Predicted differences between mean offspring *EPSPS* copy number and parental midpoint *EPSPS* copy number under Mendelian inheritance. Differences are consistent with random sampling error (∼equal differences above and below zero, dashed horizontal line) and independent of parental midpoint.. D) Observed differences between parental midpoint EPSPS copy number and mean offspring *EPSPS* copy number for *A. palmeri* (n= 39 crosses). In data panels B) and D) blue points=controlled experimental crosses, coral-red points=highly isolated experimental crosses.

## 2. Methods

### 2.1. Heritability of gene copy number

#### 2.1.1. Copy number as genomic phenotype

Here we apply a classic approach from quantitative genetics to estimate *h*^2^ (narrow-sense heritability), the proportion of phenotypic variance that is explained by (additive) genetic variance. The slope of the linear regression of the ‘midparent value’ – mean of the mother and father trait values – on the mean phenotype of the offspring estimates narrow-sense heritability (*h*^2^); this is equivalent to the proportion of variation in gene copy number that can be explained by an additive model V_A_ / V_T_ [50]. Heritability has an upper-bound slope of 1 because environmental effects and non-additive genetic effects are included in the denominator (V_T_). In this experiment, plants were grown in a relatively benign glasshouse environment and glyphosate stress was not directly applied, thus minimizing environmental effects on estimates of copy number. Here we investigate the heritability of a genomic trait rather than an organismal one, which we denote as *h*_G_^2^, by directly estimating copy number of a specific resistance gene per diploid genome. Under a purely Mendelian model of segregation and independent assortment, parents that vary in chromosomal copy number would yield offspring copy number intermediate to parents. Consistent with this Mendelian prediction, examples of copy number variants exhibiting the expected high (∼1) *h*_G_^2^, and with offspring exhibiting mean copy number intermediate to parentals, have been detected in the human genome [40]. Departures from this expected high (∼1) *h*_G_^2^ for a copy number variant suggest non-Mendelian inheritance.

#### 2.1.2. Plant material and growth conditions

*Amaranthus palmeri* seeds used in this study were sourced from open-pollinated agricultural fields in Georgia, North Carolina, and Illinois USA. Seeds from populations GA-IL and WO-GA, in which individuals exhibit a wide distribution of *EPSPS* copy number and report high mean phenotypic glyphosate resistance and amplified *EPSPS* gene copy number [49], were used for the initial heritability study. Seeds were germinated in 3.81 cm plug trays using Sungro Professional Growing Mix #1 in a controlled greenhouse environment (22-30°C, 15:9h day/night cycle) at [redacted] Phytotron. Seedlings were transplanted into 15.24 cm pots 3-5 days post-germination, and fertilized once with 20-20-20 nitrogen-phosphorous-potassium (50 ppm) one week after transplanting, and once more during inflorescence development. n.b. Specific history of glyphosate use in populations of origin is not known, although past glyphosate use is expected; during our experiments no glyphosate was applied.

#### 2.1.3. *DNA extraction and ddPCR estimation of* EPSPS *copy number*

Fresh leaf tissue (2-3cm in length) from the third and/or fourth leaves from the shoot apical meristem were collected two weeks post-transplant. All fresh samples were flash-frozen in liquid nitrogen, and subsequently stored at -80°C for DNA extraction. Genomic DNA was extracted using a modified CTAB (cetyltrimethylammonium bromide) protocol [49]. DNA purity and concentration were assessed using a DeNovix DS-11 FX+ spectrophotometer, and samples were standardized to 0.7ng/μL. *EPSPS* gene CNV was quantified using droplet digital PCR (ddPCR, Bio-Rad QX200). Each 20μL reaction included 1× ddPCR Supermix for Probes (no dUTP), 340nM forward and reverse primers for target *EPSPS* and single-copy reference gene *ALS* (acetolactate synthase), 250nM targe and reference probes (HEX and FAM-labeled respectively, Primer and Probe sequences in [49]), 0.2μL of 2U HindIII, and 5μL of DNA template. Droplets were generated using the Bio-Rad DG8 Cartridge, and amplified using a BioRad deep well thermocycler (C1000 Touch^TM^; 95°C for 10 mins; 94°C for 30 s, 56°C for 1 min, 72°C for 30 s, repeated 49x; 98°C for 10 mins; and 4°C for 10 mins). Droplets were assayed with QX200 Droplet Reader (BioRad 1863001), and the ratio of positive droplet *EPSPS*:*ALS* amplifications was analyzed using QuantaSoft/QX Manager Software (BioRad). Samples with droplet counts <10 or >10,000 for either the target or reference genes were subsequently re-diluted (0.1ng/μL to 2.0ng/μL) and re-run (following [redacted REF]). Replicability of copy number estimates using this method is high (i.e. set of 31 samples ranging in *EPSPS* copy number from one to ∼93 run in duplicate at template concentration 0.7 ng/μl; correlation between duplicates was *r*=0.99, *p*<0.0001).

#### 2.1.4. Controlled crosses by resistance gene copy number

To investigate *EPSPS* CNV inheritance, controlled crosses were performed between (full sibling) female and male plants based on *EPSPS* copy number estimates (above). Prior studies [30,39] focused on crossing single/low copy x high copy number individuals to establish whether high copy number could be transmitted to the next generation, and confirmed that offspring from such matings can carry many (up to 91) copies. Therefore, to better understand patterns of heritability, and to represent the wide variety in matings between copy number variants, we included crosses between females and males with both similar and varying parental copy number. Crosses (n= 31) ranged in parental midpoint *EPSPS* copy number from 17.5 to 92.3 (Table S1). And, crosses varied in the magnitude of difference (range 0.4 to 52.5) between maternal and paternal copy numbers. Similar numbers of crosses were performed in which maternal copy number was greater than paternal copy number (n=16 crosses), and in which paternal copy number was greater than maternal copy number (n=20 crosses). As plants approached flowering, female and male pairs were enclosed in custom-made pollination bags (PBS International, 3D.55), with an average pore size (15μm) smaller than estimated range of *A. palmeri* pollen diameter (21-38μm) [51]. In cases where anthers dehisced prior to bagging, male plants were spatially isolated for 24h, longer than the estimated length of pollen viability, prior to being paired with female plants to minimize pollen contamination among crosses [51]. Mature offspring seeds were harvested from each controlled cross. The seed count for five representative inflorescences, and their corresponding inflorescence lengths (mm) were used to estimate total offspring seed count per cross. Offspring (mean n=25/cross; range 8-36) were germinated, grown, and *EPSPS* copy number estimated using the same methods as detailed above for parents.

Additional crosses were performed, motivated by the broad range of offspring copy number observed in the first experiment. Our goal was to assess whether pollen contamination between experimental crosses conducted concurrently is a likely contributor to observed variation in offspring copy number. Sex-specific markers [52,53] were used to identify females and males, allowing female-male pairs to be isolated in bags prior to plants becoming reproductive. We further eliminated the possibility of pollen contamination by restricting the copy number variation of reproductive *A. palmeri* grown in our experimental facilities: First, ‘control’ crosses were created March to mid-April, 2022) to confirm that crosses between single *EPSPS* copy number individuals yield single-copy offspring. We used seed from a population that is predominantly composed of single *EPSPS* copy number individuals (EF-IL), confirmed via ddPCR that each parent carried a single copy EPSPS (Table S1), and then crossed three female-male pairs. Note that no other *A. palmeri* was reproductive in the growth facility during the reproductive period of these 1×1 crosses Then, seed from GA-IL was again used to experimentally cross six additional pairs (late April to May, 2022), with *EPSPS* copy number parental midpoints ranging from 14.6 to 91.0: Two lower copy crosses (parental midpoints 14 and 18) were isolated in one greenhouse, three higher copy crosses (parent midpoints 52-91) in another separate greenhouse, and one mixed (parental copy numbers 21 x 77) in a third greenhouse to further minimize opportunity for contamination between lower and high copy number crosses. As in the original crossing experiment, mature seed were collected and an average of 27 offspring per cross were grown and assayed for *EPSPS* copy number.

#### 2.1.5 Data analysis: EPSPS copy number heritability

All data analysis here and in section 2.2.2 (below) was performed in R (v. 4.2.1) and RStudio (v. 2024.04.2+764). We examined the relation between parental midpoint (mean) and offspring mean *EPSPS* copy number as follows. First, we applied a linear (lm) model to estimate the slope of the regression between parental midpoint and offspring mean. The residuals of this model did not deviate from normality (Shapiro-Wilk normality test: *W* = 0.98, *p* = 0.62), and thus a gaussian distribution was used for the subsequent threshold analysis. To test for non-linear heritability, we applied a linear quadratic model, and a segmented threshold model (library ‘chngpt’) [54] to identify any potential variation in the slope of the relation between parental midpoint and offspring mean copy number across the dynamic range of *EPSPS* copy number. The fit of these three models were compared with AIC (Akaike Information Criteria) and BIC (Bayesian Information Criterion). n.b. Modeling the relation using a generalized linear model with Poisson distribution yielded worst fit; including both linear and quadratic terms improved fit, but nonetheless much worse fit than models presented below (section 3.1). We ran each of these models using data from all crosses (n=39), and with 1×1 crosses excluded (n=36).

We tested the Mendelian prediction that differences between offspring and parents should be ∼equally positive and negative by examining the relation of difference between parental midpoint and offspring mean copy number against parental midpoint; this relation was similarly examined with linear and threshold models. And, we tested whether crosses with larger differences between parental copy numbers are associated with high s.d. of offspring copy number. Finally, we compared linear relations between offspring copy number and each of maternal and paternal copy number to test whether offspring copy number is biased by either parent.

### 2.2. Estimating intra-individual variation in *EPSPS* copy number

#### 2.2.1 Intra-individual sampling

Seeds were germinated from four *Amaranthus palmeri* populations (GA-IL, WO-GA as above, plus WA-NC, RA-NC) (Table S1 in [redacted]). Two of these populations intentionally overlap with the populations utilized in the heritability study above. We studied populations from three eastern states; although we had no *a priori* prediction that intra-individual variation in *EPSPS* would vary geographically we included samples from three geographic regions to increase generality of results. A total of 117 plants from four populations (22-39 plants per population) were studied. Seeds were germinated in the same conditions as above, but with 12/12h light/dark photoperiod, transplanted and grown as above.

From each individual plant (n=117), a series of leaf tissue was collected from a standardized position (3^rd^ or 4^th^ leaf from the apical meristem), over time, to capture variation in copy number among leaf tissues, and any potential temporal variation in leaf tissue copy number as plants develop. The first leaf sample was collected from juvenile plants at 10-14 days post-transplant. Following the first sampling, leaf tissue was collected at 10-day intervals, for a total of five temporal replicates per plant (except for four plants, for which only four temporal replicates were successfully collected; total of 881 leaf tissue samples from the 117 plants). At each temporal sampling point, biological replicates were randomly chosen for ∼20% of samples (109 leaf pairs, for 22-25 plants per sampling point) by dividing one leaf approximately in half. Inflorescence tissue was also sampled in biological replicates (1cm per replicate) from the primary inflorescence once it reached a minimum length of 4-5cm for 29 plants. All fresh tissues sampled were placed in 1.5 mL microcentrifuge tubes on ice during sample collection, and then immersed in liquid nitrogen to flash freeze samples before storage at -80°C. Each biological replicate underwent DNA extraction and analysis of copy number independently as detailed above (see 2.1.3).

#### 2.2.2. Data analysis: intra-individual variation in EPSPS copy number

Using the temporal replicate estimates of *EPSPS* copy number, we first examined the relation between mean individual *EPSPS* copy number and intra-individual variance relation relative to a Poisson expectation (E(x)=Var(x)). Residuals were calculated as the difference between observed variance in intra-individual copy number and expected variance (individual mean copy number), and standardized by dividing by s.d.(residuals). We then examined the relation between standardized residuals and mean copy number using a linear model. Due to overdispersion of *EPSPS* copy number, we subsequently used Quasi-Poisson generalized linear models, which fits a dispersion parameter to account for additional variance, to examine variation in *EPSPS* copy number with the following models and explanatory effects: 1) Overall copy number variation was tested with individual, population, temporal replicate (leaf tissue) nested within individual and biological replicate nested with temporal replicate and individual (leaf tissue); 2) Directionality of temporal replicates were tested using individual and temporal replicate nested with individual to allow for estimation of slope and significance for each individual; Bonferroni-corrected α=0.05/117=0.0004. Finally, tissue type (leaf, inflorescence) was tested for a small subset of individuals (n=11 individuals, ranging in mean copy number from 1 to 81) for which both a pair of leaf and inflorescence biological replicates were assayed; tissue type * biological replicate nested within individual (Poisson was sufficient).

Finally, to investigate whether offspring from controlled crosses with copy number (2.1.3 and 2.1.4 above) greater than the sum of parental copy numbers could be the result of intra-individual copy number variation, we applied the following analysis. For each parent we estimated the maximum range of intra-individual variation by averaging across up to 5 of the closest mean copy number estimates to parental copy number; the maximum intra-individual range of the two parents was averaged. The maximum range divided by two (because here interested in the potential for positive bias) was added to each of the parents as an upper estimate of parental copy number. Parental sum copy number was re-calculated based on these upper estimates, and the proportion of offspring (if any) above this more stringent limit calculated.

## 3. Results

### 3.1 *Non-linear inheritance of* EPSPS *copy number*

Overall, we detected significant narrow-sense heritability (LM: *ϐ/h_G_^2^*(± s.e.)=0.66±0.08, *t*=7.76, *p*<0.0001; Fig. S1) based on the linear regression between parental midpoint and mean offspring *EPSPS* copy number. A significant quadratic relation (LMquadratic: *F*(2,36)=85.76, *R^2^_adj_*=0.82, *p*<0.0001; Fig. S1) was also observed, reflecting a peak in offspring copy number at moderate parental midpoint copy number. A threshold model identified a changepoint in the slope at 33.2 (95% CI=28.1 to 57.0; Fig. S2A) copies (Fig. 1B): crosses with lower copy number have high heritability (*h_G_^2^*(± s.e.)=1.60±0.25), *p*=<0.0001), whereas heritability was much lower (*h_G_^2^*(± s.e.)=0.22±0.25, *p*<0.0001) at high copy number. Of the non-linear models, the quadratic model is the best fit (AIC=277.64, BIC=284.29), although non-linear threshold model fit (AIC=280.57, BIC=288.88) is similar to the quadratic model, and both non-linear models exhibit substantially better fit than the linear model (AIC=306.26, BIC=311.25). Excluding the three single-copy crosses yields similar results (see Supplement A) with a threshold changepoint of 48.8 copies.

The difference between mean offspring copy number and parental midpoint copy number exhibits a pattern of strong directionality across parental midpoint (Fig. 1D), which shifts from positive to negative across the dynamic range of *EPSPS* copy number (Fig. 1D, Fig. 2A and Fig. S2B). This shift in directionality was observed across the initial set of 30 crosses (Fig. 1D, blue points) and data points from highly isolated crosses (Fig. 1D coral red points) are consistent with this pattern. Crosses with parental midpoint copy numbers below 33.2 copies (Threshold model: segment 1, β=0.60±0.25, *p*<0.02), all consistently yielded offspring with more copies on average, than the parental midpoint copy number, with a maximum mean increase of 26.8 copies. Above 33.2 copies, the slope reversed from positive to negative (Fig. 1D), and the first mean decline in offspring copy number was observed at parental midpoint ∼50.4. All eight of the highest parental midpoint crosses (parental midpoints >60.3+) exhibited a mean decrease in *EPSPS* copy number for offspring relative to parental midpoint. And, the decline in offspring copy number increased in magnitude with increasing parental midpoint copy number (Threshold model: segment 2 β=-0.78±0.20, *p*<0.0001), with a maximum mean decrease of 37.7 copies which was observed in a highly isolated cross.

**Figure 2.**
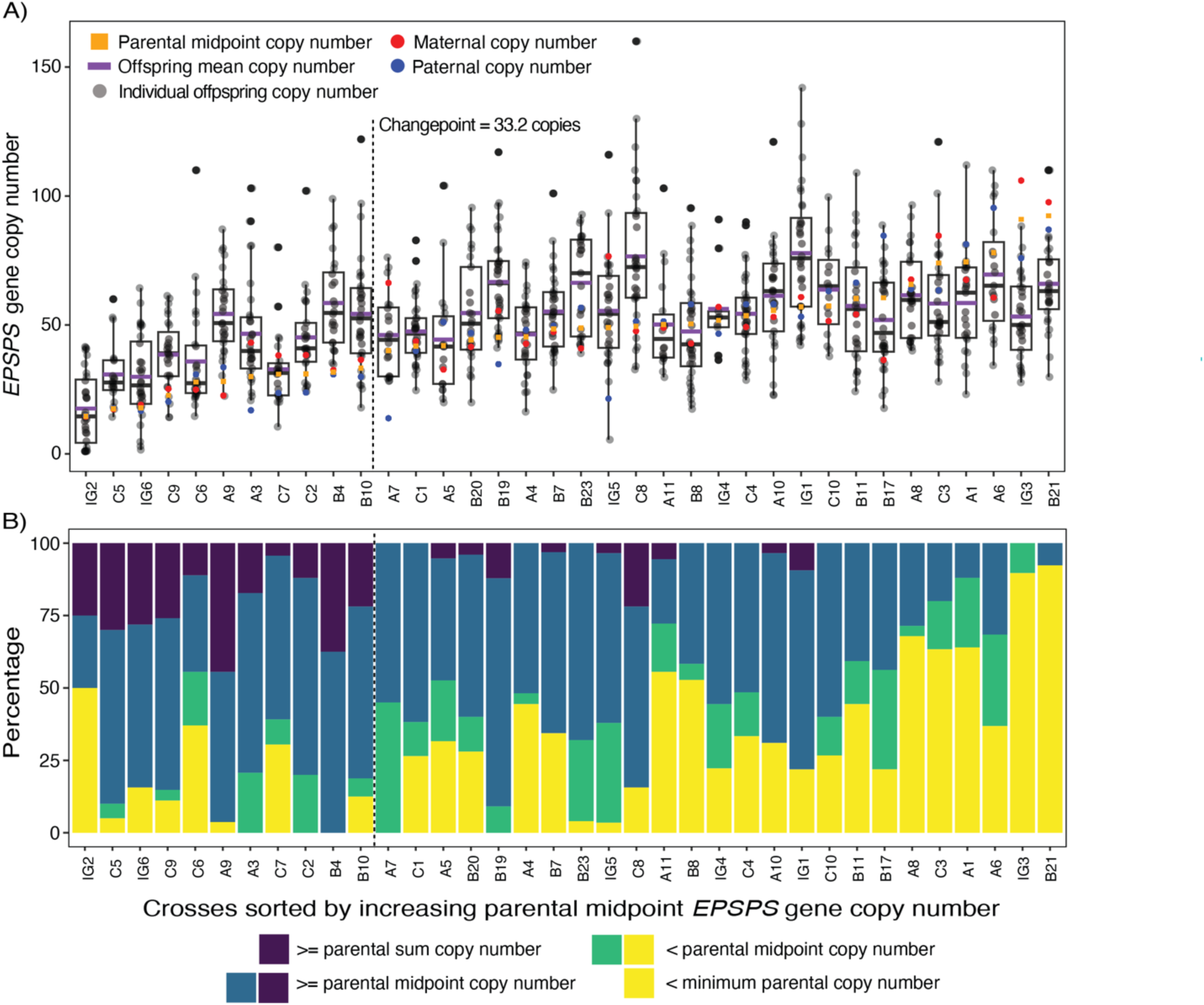
Variation in *A. palmeri* offspring *EPSPS* copy number relative to parental *EPSPS* copy number. A) Boxplots of *EPSPS* copy number variation for each experimental cross. Orange-yellow squares=parental midpoint copy number; Purple lines=offspring mean copy number; Gray points=individual offspring copy number; Red circles=maternal copy number, Blue circles=paternal copy number, boxplot median line=offspring median. B) Bar plot of percent of offspring exhibiting copy number less than parental midpoint copy number (green and yellow) and greater than or equal to parental midpoint copy number (blue and dark purple). Dark purple=percent of offspring exhibiting copy number greater than the sum of parental copy numbers. Yellow=percent of offspring exhibiting copy number less than the lowest parental copy number. Box and bar plots in A) and B) are sorted from left to right by increasing parental midpoint. The occurrence of the changepoint (parental midpoint 33.2 gene copies) identified in Figs. 1B and D relative to the sorted crosses is shown with vertical dotted line in both panels. Each stack of bars in panel B corresponds to the cross displayed directly above in boxplot A.

The observed shift in directionality also applies to offspring exhibiting extreme high and low copy numbers, relative to parents. Below the changepoint (up to 33.2 copies, dotted line on Fig. 2B), on average 23% of offspring exhibited copy number greater than the sum of both parental copy numbers. In contrast, above the changepoint on average 2.7% of offspring exhibited copy number greater than both parental copy numbers (Fig. 2B, dark purple bars). And, below the changepoint on average 15% offspring exhibited copy number lower than the lowest copy number parent, whereas above the changepoint on average 36% did (Fig. 2B, yellow bars).

We tested whether larger differences between individual parental copy numbers result in more dispersion in the distribution of resulting offspring copy numbers, and found no significant relation between the difference in parental copy numbers and the standard deviation of offspring copy number (lm: β=0.13, *t*=1.88, *p*=0.07; Fig. S3). No evidence of large sex-specific effects were detected; the relation between parent and offspring copy number is similar when based on maternal copy number (β=0.64, *t*=7.17, *p*<0.0001) or paternal (β=0.49, *t*=5.84, *p*<0.0001) copy number.

### 3.2. EPSPS copy number variation: intra- and inter-individual sources

Intra-individual temporal replicates facilitated testing of the relation between individual mean and individual variance for estimates of *EPSPS* copy number. Overall, we observed a clear increase in copy number variation among temporal replicates as individual mean copy number increases, with variation ∼symmetrical relative to 1:1 (Fig. 3A). The Poisson expectation for count data is that variance of intra-individual samples will increase due to increasing mean count alone (Fig. S4A, red line). As mean copy number increases, individuals exhibited variance of temporal replicates far greater than expected due to the Poisson expectation alone (Fig S4A). A significant positive (LM: *R*^2^ = 0.33, *t*=7.57, β=0.03 *p* < 0.0001) relationship between mean copy number and standardized residuals verified a significant increase in the deviation from the Poisson variance expectation as mean copy number increases, and indicated overdispersion relative to the Poisson expectation (Fig. S4B), consistent with the overall estimate of dispersion χπ=1.98 (if χπ>1 Poisson underestimates variance).

**Figure 3.**
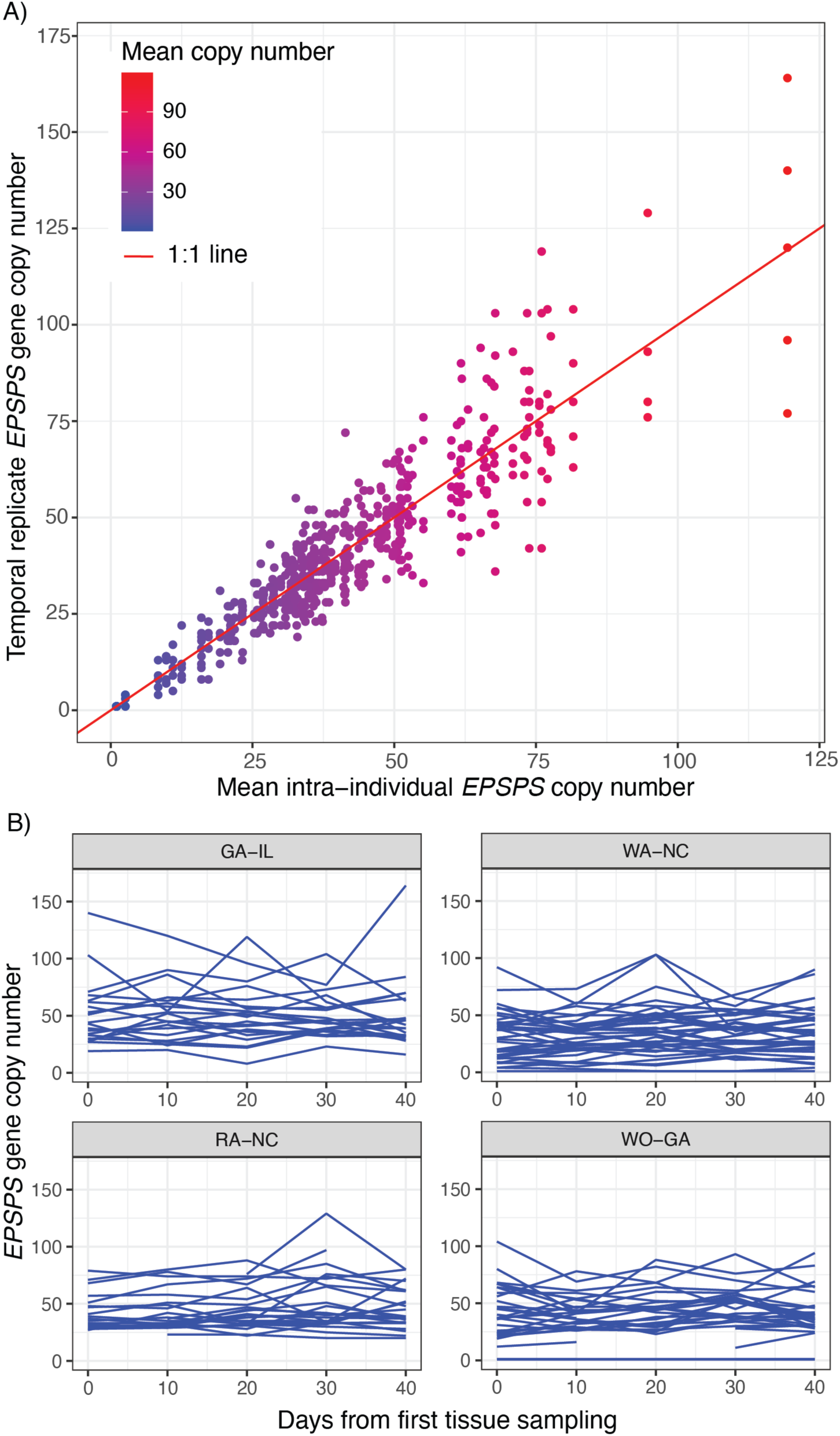
Intra-individual variation in *EPSPS* copy number in *Amaranthus palmeri*. A) Scatterplot of individual (leaf tissue) estimates of *EPSPS* copy number by mean intra-individual copy number; intra-individual variation increases with mean copy number. Red line=1:1. B) Temporal intra-individual variation based on five leaf tissue samples for four *A. palmeri* populations (GA-IL, RA-NC, WA-NC, WO-GA). Each blue line joins (up to) five estimates of *EPSPS* copy number from leaf tissue collected at 10-day intervals within an individual *A. palmeri* plant; all temporal relations were non-significant following Bonferroni correction.

Using a Quasi-Poisson specified GLM model to account for the overdispersion of copy number, we found that the majority of variation in *EPSPS* copy number occurs among individual plants (GLM: *F*_66,155_=70.93, *p*<0.0001), with lower-magnitude variation among populations also detected (GLM: *F*_3,221_=42.17, *p*<0.0001). But, two sources of intra-individual variation were also significant: Biological replicate, although significant, explained the lowest amount of variation in *EPSPS* copy number (GLM: F_70,55_=1.76, *p*=0.02), with temporal replicates explaining more variation (GLM: *F*_30,125_=6.99, *p*<0.0001).

*EPSPS* copy number estimates of biological leaf tissue replicates (n = 109 pairs) differ by 5.83 (s.d. 6.97) copies on average, with differences ranging from 0 to 53 copies. Difference between biological leaf tissue replicates was only statistically significant for one individual from WO-GA following Bonferroni correction (α=0.05/70=0.0007). And, tissue type (leaf vs. inflorescence) did not significantly affect variation in biological replicates (GLM: tissue type x individual:biological replicate *z* = 0.05, *p* = 0.95). Temporal replicates of leaf tissue on average exhibited maximum pairwise difference in copy number of 20.64 copies (s.d. 14.50; range 0-87 copies). We tested whether variation observed across (five) temporal replicates per individual (n=117) exhibit directionality (ie: significant increase or decrease), and after Bonferroni correction (α=0.0004) no individuals exhibited significant directional change in copy number across temporal replicates (minimum *p*-value observed is 0.0012) (Fig. 3B; blue lines).

## 4. Discussion

Gene copies carried by extrachromosomal DNA are predicted to be less stable than chromosome-integrated tandem repeats, due to the dynamic association with chromosomes via tethering during mitosis and meiosis [30,55]. Here we show that copy number of resistance gene *EPSPS* is unstable within tissues and among offspring, and that the level of instability is predictable by mean copy number. Controlled experimental crossing revealed a non-linear pattern of inheritance (Fig. 1). A threshold suggests high heritability of copy number variation at lower copy number, and decline in heritability at higher copy number. This shift in heritability was due to the fact that offspring mean copy number was consistently maintained or increased at lower mean parental copy number (Fig. 2), but declined steadily across the highest parental copy numbers. Similarly, the stability of copy number variation within individuals decreases with mean plant copy number (Fig. 3). Here we discuss these deviations from Mendelian expectation in the context of *A. palmeri’s* extrachromosomal circular DNA structures as a mechanism driving copy number variation and herbicide resistance. We evaluate the potential for genomic instability to contribute to the observed pattern of *EPSPS* inheritance, and potential limits to evolutionary change.

### 4.1 Heritability of an extrachromosomal copy number variant: deviations from Mendelian expectations

As a function of additive genes and environmental influences, narrow-sense heritability for both organismal and genomic traits are expected to vary among estimates [36]. Overall, our estimate of *h*_G_^2^=0.66, is higher than a previous estimate *h*_G_^2^=0.29 [39]; this difference may be owing to the previous study being based mostly on crosses between single/low copy x high copy individuals, whereas our study deliberately included crosses between parents with more similar copy numbers. Yet, these estimates of *h*_G_^2^ are lower than *h*_G_^2^∼1 [40] for human chromosomal gene copy number variation. In contrast to previous studies that report a single heritability value we found the pattern of inheritance varies across the dynamic range of copy number. Heritability (*h*_G_^2^) of *EPSPS* copy number is high (1.60) below the threshold of ∼33 gene copies, and above heritability’s theoretical limit of one [56]. Above the changepoint (∼33 copies), heritability is substantially lower (*h_G_^2^*=0.22). ^Thus, just as heritability of an^ organismal phenotype can vary between sexes or populations [36,57], this genomic phenotype – *EPSPS* copy number – varies by genotype. This change in heritability can be explained as differences in additivity, which determines the numerator of *h^2^,* differences in environmental influences which affect the denominator, or both.

Random measurement error should be symmetric (Fig. 1C) but instead we found a positive bias in differences between offspring and parents across the lower range of copy number, and a negative bias across the upper range (Fig. 1D). The consistency of offspring mean increases in copy number at lower copy number, and offspring mean losses at higher copy number, suggest this directionality cannot be explained by sampling error alone (Fig. 1D). Pollen contamination could cause an increase in offspring copy number in low copy crosses, and a decrease in high copy crosses. However, the consistency of results from highly isolated crosses (Fig. 1D, coral red points) and the original crosses rules out this possibility.

The heritability above one at low copy number suggests *EPSPS* copy number is stable, and possibly associated with a capacity for increase through meiosis and sexual reproduction. Previous visualization of eccDNA suggests that the lower part of the copy number range (1 to at least 12 copies) can be chromosomal (possibly tandem) copies, rather than positioned on extrachromosomal DNA [30]. Thus, this low range of copy numbers are predicted to be consistent with Mendelian expectations, and indeed our 1×1 copy crosses yielded all but one (see Supplement B) single copy progeny. Copy numbers above 12 are increasingly likely to reside on extrachromosomal DNA [30], and parental midpoints up to ∼50 copies result in stable transmission of these copies to offspring. The difference between mean offspring and parental midpoint copy number consistently ranges between 2 and 27 copies across lower copy numbers (segment 1, Fig. 1B), raising the question of whether consistent positive bias of mean offspring copy number is the result of a molecular mechanism for the further replication of the *EPSPS* extrachromosomal replicon.

The positive bias in offspring copy number is expected for the lowest range of copy numbers due to the constraint of requiring at least one copy of *EPSPS*, but this constraint lessens with increasing copy number. Instead, the positive bias persists up to (parental midpoint) ∼50 copies − this is consistent with Koo et al.’s [30] observation that eccDNA structures randomly associate with chromosomes through the pachytene, diplotene, diakinesis, and metaphase stages of meiosis. Thus, there is a mechanism by which an offspring could inherit multiple eccDNAs/copies from each parent in addition to chromosomal gene copies. Up to the changepoint of 33 copies, 77% of individual offspring exhibited *EPSPS* copy number estimates greater than the parental midpoint and 23% of offspring exhibited copy number greater than the sum of both parents. In fact, the 33-copy changepoint coincides with the peak of the increase in copy number of offspring relative to parents: for cross B10 between parents with 37 and 30 copies the maximum offspring value was ∼122 copies, double the sum of both parents. In contrast, for a lower copy number cross between 14 and 14 copies, the maximum offspring value was ∼13 copies greater than the parental sum. The large gains in copy number drop precipitously below the changepoint (mean offspring > parental sum above the changepoint is 3%). In contrast to inherited copies, intra-individual temporal replicates suggest that there is no general positive increase in copy number through plant development (Fig. 3B).

Intra-individual variation in copy number did not account for the range of variation observed in offspring (see 2.2.2 and Supplement C). For all but one cross, up to 19% of the highest copy number offspring are higher than the sum of estimated variation among tissues of the parents. Thus overall, mechanisms responsible for intra-individual variation likely contribute to the range of copy number gains observed in offspring, but are insufficient to explain the consistently positive bias of offspring relative to parents, or the most extreme high copy number offspring that exceed both parents. Specifically, the extrachromosomal *EPSPS* replicon, once introduced to a population, can generate a wide range of copy numbers within a few generations. In principal, this is equivalent to a rapid increase in the standing genetic variation of a quantitative trait, which increases the rate of adaptive evolutionary response to natural selection [50]. To better understand the mechanisms of these rapid gains in copy number, further work is needed to characterize molecular mechanisms for the eccDNA replicon through meiosis and sexual reproduction.

In contrast to low copy number, a decline in heritability and a reversal to copy number loss was observed above ∼33 copies of the *EPSPS* gene. The significant negative slope above the changepoint (segment 2, Fig. 1D) suggests that the magnitude of copy number loss in offspring increases with copy number. One general hypothesis for this loss of heritability is that as mean individual copy number increases, some aspect(s) of the genomic environment increasingly interferes with the maintenance and/or transmission of extrachromosomal copies. The structural polymorphism of eccDNAs characterized by Koo *et al*. is consistent with extrachromosomal structures being prone to structural damage (e.g. ∼50% intact circular vs. ∼50% linear and irregular forms). Extrachromosomal DNA polymorphism was assayed for an individual carrying ∼80 gene copies [30]; it would be of future interest to test whether a greater proportion (>50%) of eccDNAs in the range of 15 to 33 copies are circular, being putatively the most complete and stable eccDNAs. This would test whether stability of circular forms plays a role in the stable inheritance and positive copy number bias observed here over lower copy number range. Further, the stronger decline at high copy number suggests a potential for influence of the spatial arrangement and density of eccDNAs. For example, perhaps high density of eccDNAs contributes to a larger proportion of damage, and lowers transmission of gene copies during meiosis.

The magnitude of mosaicism [58,59] of *EPSPS* copy number [30,39] is shown here to increase in variability, becoming over-dispersed as mean copy number increases (Fig. 3A, Fig. S4). This suggests that copy number of gametes could be associated with high variance, yet we observe the striking negative bias of offspring copy number. This suggests that copy number could have direct genetic effects on gamete performance. Copy number can be associated with either positive or negative effects on fertility: female gametes in potato were found to benefit from increased copy number in a linkage group [60], whereas human male infertility has been associated with increased gene copy numbers on the Y-chromosome [61]. In plants, variation in pollen tube growth rate can be a significant component of gametic competition [62,63]; low heritability of high copy number could be the result of high copy number pollen tubes being outcompeted by lower copy number pollen tubes. A first step toward testing this hypothesis would be to quantify the variance in copy number among pollen grains, followed by comparison of pollen tube success by copy number.

### 4.2 Evolution of copy number and genomic plasticity

High copy number individuals exhibit the most consistently high phenotypic glyphosate resistance, but the gains in resistance per copy are small from 16-160 copies [49]. Moreover, the signature of cost of resistance / high *EPSPS* copy number detected in other glyphosate-resistant plants [64–66]– a negative relation between copy number and fitness-related traits in the absence of glyphosate – is often not detected in *A. palmeri* [67–69]. Here we detect no relation between seed count of crosses and parental midpoint copy number (lm: *t*=1.43, *p*=0.16). The only evidence of a cost was detected by a field assessment and found a fitness penalty (lower seed number and biomass) associated with *EPSPS* copy number >2 copies in *A. palmeri* [69]. Therefore, given the modest costs associated with high copy number but the consistent highest resistance conferred, it has seemed paradoxical that high copy number individuals have only been detected in agricultural populations at very low frequencies [49]. One possibility is that more time is required and directional selection, if glyphosate selection continues, will further increase mean *EPSPS* copy number in these agricultural populations [27,67]. To date, the highest population mean *EPSPS* copy number detected is ∼58 [49], which is near the upper 95% confidence interval (∼57 copies) of the threshold copies detected here. In addition, populations with higher mean CN exhibited lower standard deviation [49], consistent with stabilizing selection and the possibility that population *EPSPS* copy number is already near a stable evolutionary peak. Our findings here suggest that declining heritability could be the primary factor limiting the evolution and maintenance of high *EPSPS* copy number in *A. palmeri*.

Extrachromosomal gene copy number variation has been proposed as a form of genomic plasticity [30,31], with the potential for this plasticity to play a role in adaptive evolution [43]. And, a connection to McClintock’s “sensing mechanism … when experiencing unfavourable conditions to alert the cell to imminent danger”[4] resulting in a set response to alleviate stress has been suggested [30,31]. This hypothesis would suggest that formation of circular extrachromosomal DNA is due to environmentally-induced genomic stress. McClintock proposed, based on studies in maize (*Zea mays*), stress-induced DNA breaks could yield DNA circularization as blunt ends are joined [4,70], followed by possible ongoing replication of eccDNAs in response to sustained genome stressors (e.g. glyphosate or other). McClintock also noted further duplication occurring within circular extrachromosomal DNA structures once they established [70]. Here, we recorded offspring with copy number greater than the sum of both parents (Fig. 2B, dark purple bars), and up to double the sum of parental copy numbers. Such increases in copy number are consistent with a putative self-replication mechanism, supported by the fact that the *A. palmeri EPSPS* replicon carries several genetic elements involved in DNA replication [31]. Autonomous self-replication has been documented in *Drosophila* eccDNA (e.g. rolling circle replication [71]), and occurs in the majority of yeast eccDNAs [43]. Moreover, in yeast an eccDNA was detected in preparations with low-nutrition media, but not detected in nutritionally-complete media [43], further supporting a role of environmental stress in eccDNA creation. Here we show that although copy number is more variable in high copy number individuals, over plant development copy number is not directional (Fig. 3B). Future studies of intra-individual variation and heritability under glyphosate and other stresses would be valuable: does glyphosate stress increase the production of eccDNAs / gene copies, furthering the positive bias in copy number observed in offspring here at lower copy numbers, and/or ameliorating the strong loss of copies toward the upper range of copy number? Or, is the genomic constraint on the maintenance of high copy number observed here observed under all environmental conditions?

### 4.3 Conclusion

Despite significant deviations from Mendelian expectations, the heritability of extrachromosomal copy number follows a clear pattern characterized by a shift in directionality. The lack of temporal directionality among tissues of the same individual shows that copies are not generally lost during mitosis, but rather the decline in offspring copy number relative to high copy number parents points to loss occurring during meiosis and sexual reproduction. Further work can explore whether copy number influences gametic fitness, how genome structure affects the integrity and transmission of extrachromosomal DNA, and the effect of environmental stress on eccDNA transmission. Overall, diminished heritability and the shift to directional loss of copy number described here suggest a potential constraint on the evolution of herbicide resistance via increased *EPSPS* copy number.

