## Supplements for "Departures from Mendelian inheritance of *EPSPS* gene copies in a herbicide-resistant weed"

Supplementary Materials.

Table S1. Summary of controlled experimental crosses between female and male *Amaranthus palmeri* informed by *EPSPS* gene copy number. Cross ID corresponds to labels in Figure 2. Cross IDs beginning with A, B or C are the original set of controlled crosses, and crosses beginning with I (the last nine rows) are highly isolated crosses. CN = copy number. Number of offspring (n) refers to the number of offspring assayed for *EPSPS* copy number.

| **Cross ID** | **Maternal CN** | **Paternal CN** | **Parental midpoint CN** | **Sum of parental CN** | **Number of offspring (n)** | **Mean offspring CN** | **Minimum offspring CN** | **Maximum offspring CN** |
| --- | --- | --- | --- | --- | --- | --- | --- | --- |
| A1 | 67.50 | 81.30 | 74.40 | 148.80 | 25 | 58.51 | 23.10 | 112.00 |
| A10 | 53.10 | 58.10 | 55.60 | 111.20 | 29 | 61.30 | 22.70 | 121.00 |
| A11 | 48.90 | 51.40 | 50.15 | 100.30 | 18 | 50.17 | 29.70 | 103.00 |
| A3 | 43.10 | 16.90 | 30.00 | 60.00 | 29 | 46.59 | 20.70 | 103.00 |
| A4 | 42.70 | 48.10 | 45.40 | 90.80 | 27 | 46.37 | 16.30 | 74.30 |
| A5 | 32.70 | 51.10 | 41.90 | 83.80 | 19 | 44.31 | 19.90 | 104.00 |
| A6 | 60.60 | 95.40 | 78.00 | 156.00 | 19 | 69.52 | 34.30 | 110.00 |
| A7 | 66.30 | 13.80 | 40.05 | 80.10 | 20 | 46.04 | 28.20 | 76.20 |
| A8 | 67.60 | 63.90 | 65.75 | 131.50 | 28 | 61.52 | 40.00 | 96.50 |
| A9 | 22.60 | 33.60 | 28.10 | 56.20 | 27 | 54.25 | 22.40 | 87.10 |
| B10 | 36.50 | 29.90 | 33.20 | 66.40 | 32 | 54.27 | 17.90 | 122.00 |
| B11 | 54.10 | 66.40 | 60.25 | 120.50 | 27 | 57.33 | 24.20 | 109.00 |
| B17 | 36.40 | 84.60 | 60.50 | 121.00 | 32 | 51.89 | 17.70 | 88.70 |
| B19 | 55.50 | 34.80 | 45.15 | 90.30 | 33 | 66.65 | 41.10 | 117.00 |
| B20 | 41.50 | 46.90 | 44.20 | 88.40 | 25 | 54.61 | 19.90 | 95.50 |
| B21 | 97.60 | 87.00 | 92.30 | 184.60 | 26 | 65.93 | 29.80 | 110.00 |
| B23 | 40.90 | 56.70 | 48.80 | 97.60 | 25 | 66.33 | 39.00 | 92.80 |
| B4 | 32.50 | 30.80 | 31.65 | 63.30 | 24 | 58.48 | 34.70 | 98.90 |
| B7 | 47.30 | 49.70 | 48.50 | 97.00 | 32 | 55.16 | 24.80 | 101.00 |
| B8 | 42.80 | 58.00 | 50.40 | 100.80 | 36 | 47.48 | 17.40 | 95.30 |
| C1 | 43.60 | 39.90 | 41.75 | 83.50 | 34 | 47.45 | 24.60 | 82.80 |
| C10 | 51.50 | 63.10 | 57.30 | 114.60 | 15 | 63.70 | 37.80 | 99.60 |
| C2 | 38.30 | 23.80 | 31.05 | 62.10 | 25 | 45.15 | 24.10 | 102.00 |
| C3 | 84.60 | 63.40 | 74.00 | 148.00 | 30 | 58.27 | 28.00 | 121.00 |
| C4 | 49.20 | 58.00 | 53.60 | 107.20 | 33 | 54.35 | 28.60 | 89.90 |
| C5 | 17.30 | 17.70 | 17.50 | 35.00 | 20 | 30.80 | 14.30 | 60.00 |
| C6 | 24.90 | 30.80 | 27.85 | 55.70 | 27 | 35.83 | 14.60 | 110.00 |
| C7 | 38.20 | 23.60 | 30.90 | 61.80 | 23 | 32.81 | 10.50 | 80.10 |
| C8 | 47.60 | 51.40 | 49.50 | 99.00 | 32 | 76.60 | 32.70 | 160.00 |
| C9 | 25.20 | 20.20 | 22.70 | 45.40 | 27 | 38.53 | 14.20 | 61.30 |
| IG1 | 60.80 | 53.20 | 57.00 | 114.00 | 32 | 77.82 | 42.10 | 142.00 |
| IG2 | 14.50 | 14.70 | 14.60 | 29.20 | 32 | 17.56 | 0.95 | 41.50 |
| IG3 | 75.90 | 106.00 | 90.95 | 181.90 | 29 | 53.24 | 27.70 | 88.30 |
| IG4 | 57.00 | 46.60 | 51.80 | 103.60 | 9 | 56.17 | 36.40 | 90.90 |
| IG5 | 76.60 | 21.40 | 49.00 | 98.00 | 29 | 55.42 | 5.50 | 116.00 |
| IG6 | 19.00 | 16.70 | 17.85 | 35.70 | 32 | 29.91 | 1.58 | 64.30 |
| IE1 | 0.91 | 0.96 | 0.94 | 1.87 | 32 | 1.00 | 0.86 | 1.18 |
| IE2 | 1.00 | 1.00 | 1.00 | 2.00 | 16 | 1.02 | 0.90 | 1.13 |
| IE3 | 0.99 | 1.12 | 1.05 | 2.11 | 31 | 0.99 | 0.79 | 1.14 |

Figure S1.

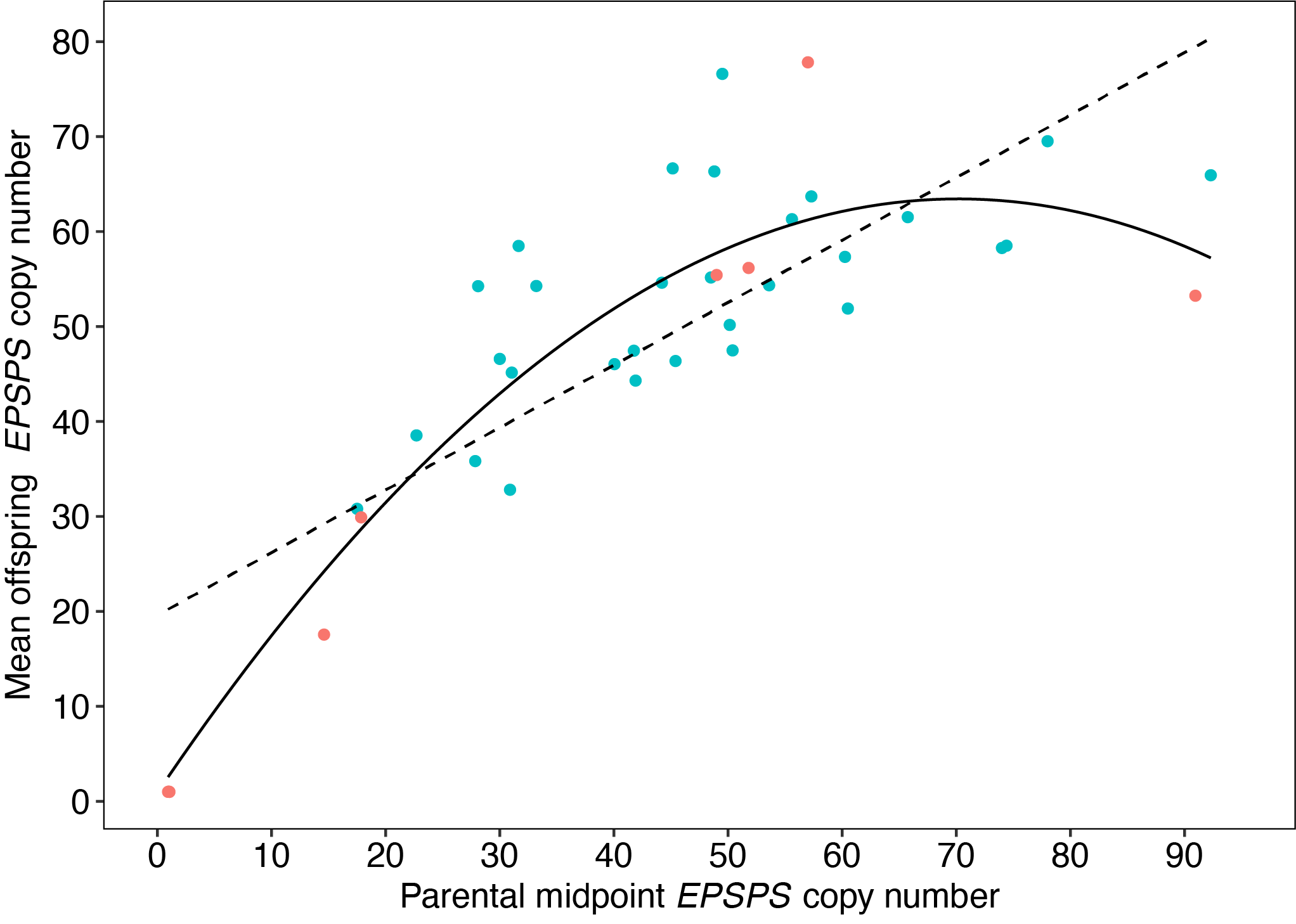

Figure S1. Scatterplot of relation between parental midpoint EPSPS copy number and mean offspring copy number in *Amaranthus* *palmeri* (n= 39 crosses). Data points displayed are the same as Fig. 1B, but with linear (dotted black line; *ϐ/h_G_^2^* = 0.66, *t*=7.76, *p*<0.0001) and linear quadratic (black line; LMquadratic: *R^2^_adj_*=0.82, *p*<0.0001, linear term *t*=9.84, *p*<0.0001; quadratic term *t*=-6.55, *p*<0.0001) fits.. Blue points=controlled experimental crosses, coral-red points=highly isolated experimental crosses.

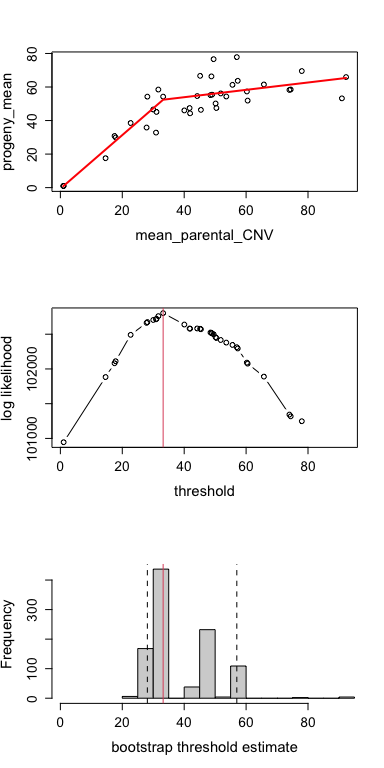
Figure S2A.

Figure S2A. Segmented threshold analysis of the relation between parental midpoint *EPSPS* copy number and mean offspring copy number in *Amaranthus* *palmeri* (n= 39 crosses). Top panel is the relation with estimated slopes for segments 1 and 2 as presented in Fig. 1B. The middle panel shows the threshold value identified (33.2 copies) by bootstrap analysis (bottom panel).

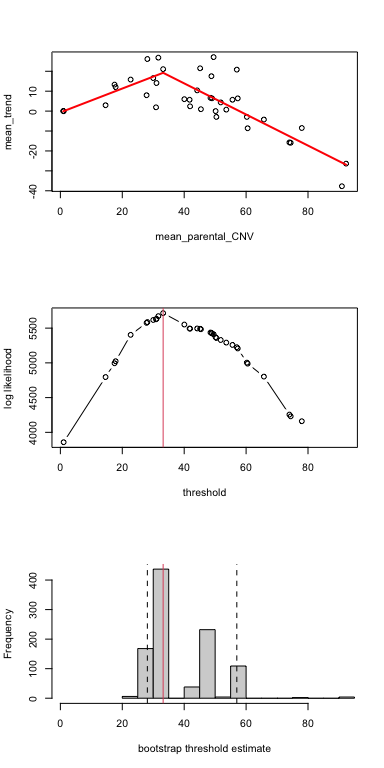
Figure S2B.

Figure S2B. Segmented threshold analysis of the relation between the difference of offspring mean copy number and parental midpoint copy number, and mean offspring copy number in *A.* *palmeri* (n= 39 crosses). Top panel is the relation with estimated slopes for segments 1 and 2 as presented in Fig. 1D. The middle panel shows the threshold value identified (33.2 copies) by bootstrap analysis (bottom panel).

Figure S3.

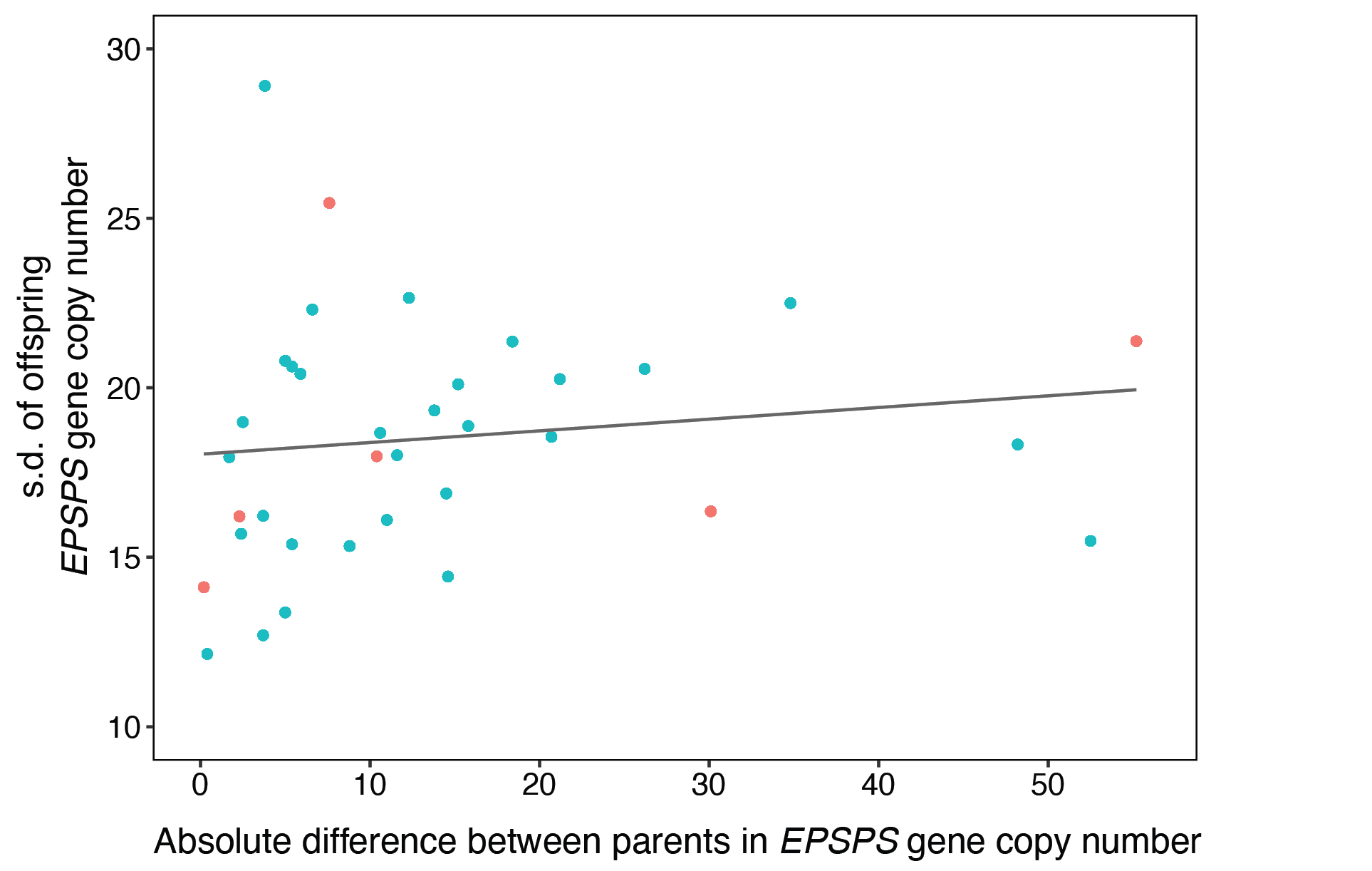

Figure S3. Relation between different in parental *EPSPS* copy number and the standard deviation of offspring EPSPS copy number for 37 experimental crosses between female and male *Amaranthus palmeri*. Blue points=controlled experimental crosses, coral-red points=highly isolated experimental crosses.

Figure S4.

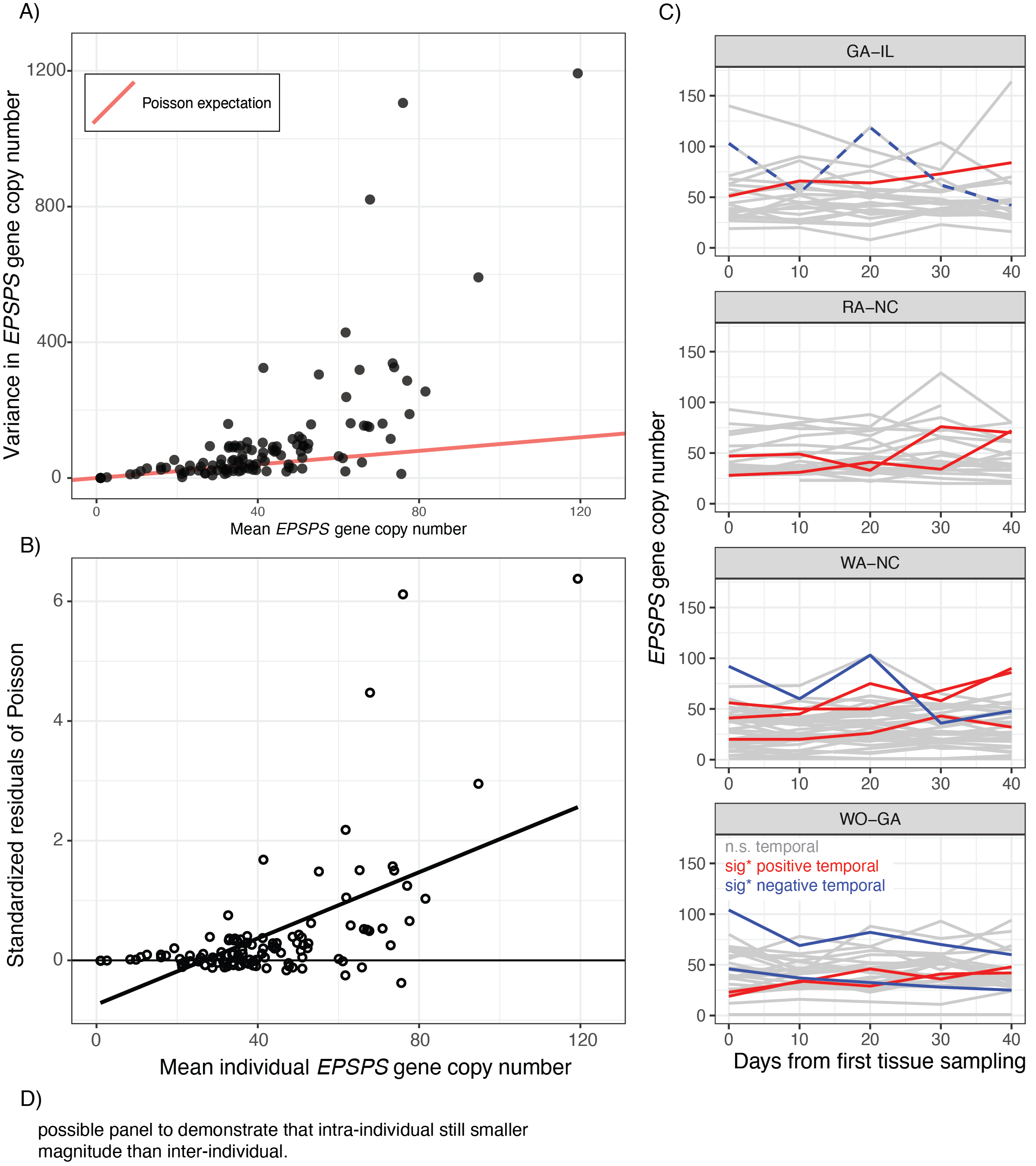

Figure S4. A) Scatterplot of mean *EPSPS* copy number estimated for five leaf tissue samples collected at 10-day intervals by the variance of *EPSPS* copy number for those five leaf tissue samples. Red line=the Poisson expectation of variance~mean for count data. B) Standardized residuals (residual/s.d. residuals) from the Poisson expectation and observed data points (from panel A) plotted against mean individual copy number across five temporal leaf samples. Black line represents the linear fit.

Supplement A.

Heritability analysis with three 1x1 (EF-IL) crosses excluded.

Linear, linear-quadratic, and threshold models:

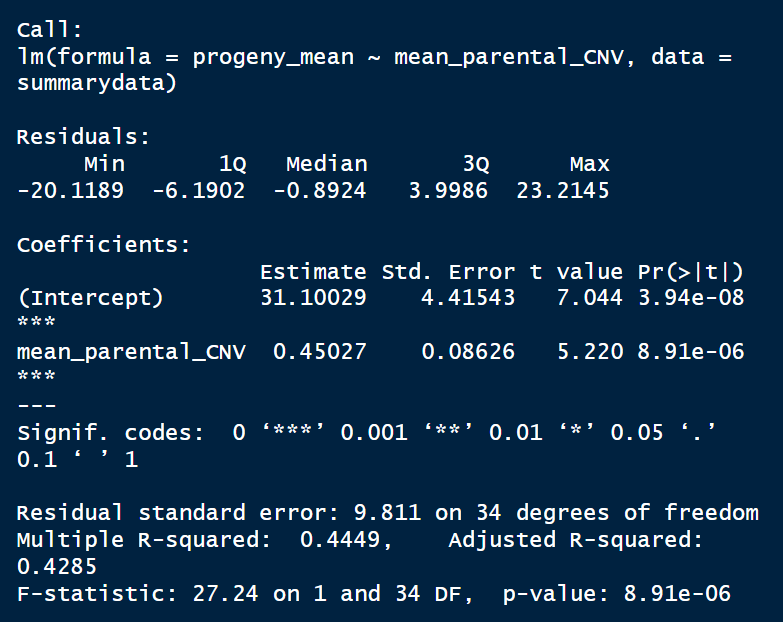

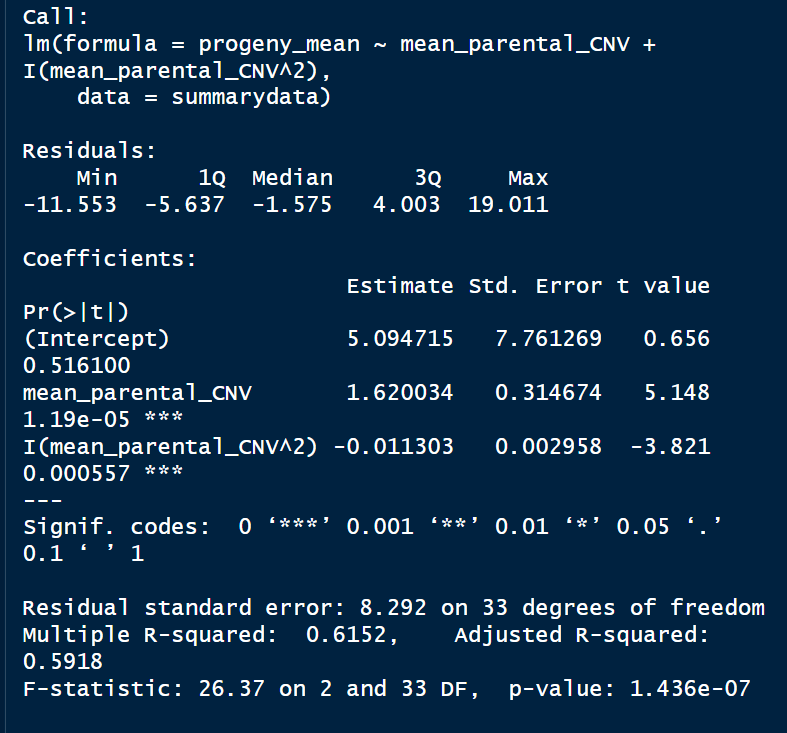

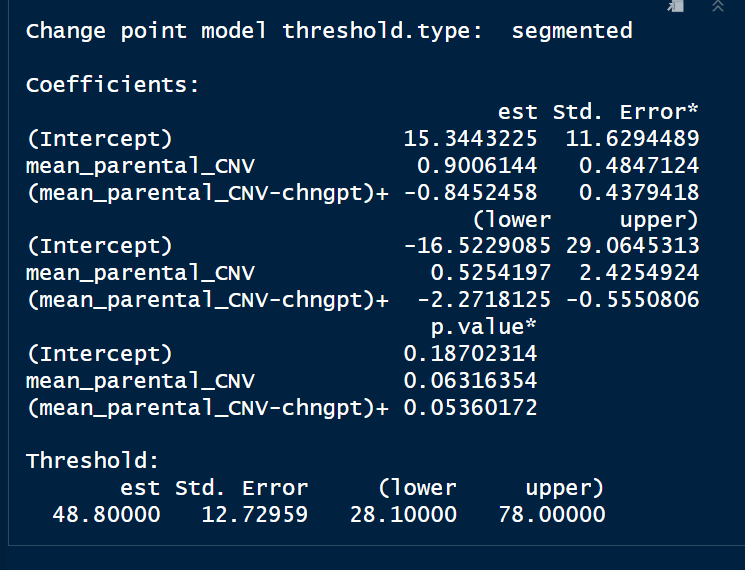

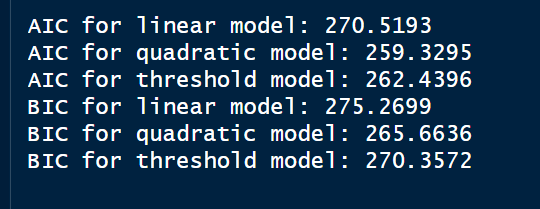

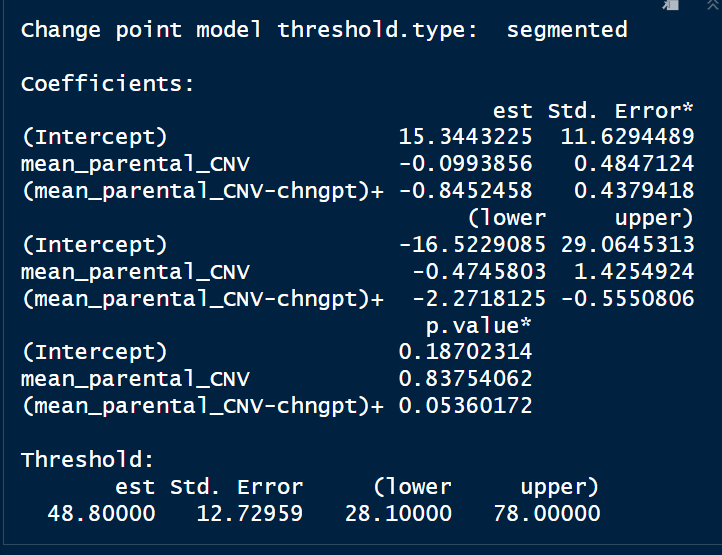

Excluding 1x1 crosses, overall, we detected a moderate estimate of heritability (LM: *ϐ/h_G_^2^* = 0.45, *t*=7.04, *p*<0.0001; Fig. S1) based on the linear regression between parental midpoint and mean offspring *EPSPS* copy number. A significant quadratic relation (LMquadratic: *F*(2,33)=26.37, *R^2^_adj_*=0.59, *p*<0.0001; Fig. S1) was also observed, reflecting a peak in offspring copy number at moderate parental midpoint copy number. A threshold model identified a changepoint in the slope at 48.80 (95% CI=28.10 to 78.00) copies (Fig. 1B): the first segment across copy numbers ~1-48 exhibits high heritability (*h_G_^2^*=0.90, *p*=0.06), but above the changepoint (parental midpoint of 48+ copies) heritability is much lower (*h_G_^2^*=0.06, *p*=0.05). Of the non-linear models, the quadratic model is the best fit (AIC=259.33, BIC=265.66), although non-linear threshold model fit (AIC=262.44, BIC=270.36) is similar to the quadratic model, and both non-linear models exhibit substantially better fit than the linear model (AIC=270.52, BIC=275.27).

Supplement B.

There are many challenges associated with the process of performing controlled crosses, especially for a wind-pollinated plant such as *A. palmeri*.

We conducted three highly isolated ‘control’ crosses, each between single copy EPSPS individuals (1x1). We estimated copy number for n=32 (IE1), n=17 (IE2), and n=32 (IE3) offspring per cross. *EPSPS* copy number estimates for all offspring from these crosses were ~1, with one exception. One offspring from cross IE3 was estimated to exhibit ~47.2 copies of *EPSPS*. We re-extracted a back-up leaf tissue sample from this offspring which yielded a similar estimate of 46.99 copies. Due to the total isolation of these crosses from other *A. palmeri*, we suspect this result is due to sample contamination of some sort. Although the crosses were executed in total isolation, the offspring were later grown in the greenhouse along with offspring from other crosses (to collect tissue for copy number estimates). Photos / records from this grow-out indicates that offspring from 1x1 cross IE3 were sown directly adjacent to offspring from high copy number cross IG5 (21x77 copies), and thus is a potential source of sample contamination through seed-hopping.

An alternative interpretation would be that single copy individuals have the capacity to vastly increase copy number through sexual reproduction. However, given the results of Koo et al. (2018) we do not expect that single copy number individuals carry the *EPSPS* eccDNA replicon, and thus we don’t expect such individuals/crosses to have the capacity for large increases in EPSPS copy number via genomic plasticity such self-replication. Thus, we consider the contamination scenario most likely, and indeed all other 80 offspring from these highly isolated crosses were estimated to carry a single copy of *EPSPS*.

Giacomini et al. (2018) report on intra-individual variation in copy number in 5 individual plants and 6 clones, which inspired the intra-individual component of our study. Their study included a single 1-copy x 1-copy cross, and reported that offspring included “29 plants with a single EPSPS copy, 1 plant with 12 EPSPS copies, and 1 plant with and 23 EPSPS copies.” The authors suggest that “progeny from GS low copy-number *A. palmeri* plants can accumulate sufficient amplified EPSPS copies to produce a GR phenotype, even in the absence of glyphosate.”, but as outlined above, contamination is a possible source that should be carefully considered. The authors also go on suggest that “progeny should be considered in relation to the within-plant somatic variation for EPSPS copy number that we also observed in this study”. In our study of intra-individual copy number variation we assayed six single-copy individuals, and for each, all estimates (5 per plant) of copy number was one. Thus, for single copy crosses we find no evidence of somatic variation in EPSPS copy number.

Due to our view that our outlier data point is likely an example of contamination, we have excluded this sample from our main analyses presented in the Results. When we do include the data point it yields a mean offspring copy number of 2.44 for cross IE3 (in contrast to estimate of offspring mean of 0.99 without this data point). Regardless of its source, inclusion of this tenuous data point does not change any of the main heritability results (as reported in section 3.1) in any substantial manner:

Linear Model:

(excluding potential contaminant) LM: *ϐ/h_G_^2^* = 0.66, *t*=7.76, *p*<0.0001

(including potential contaminant) LM: *ϐ/h_G_^2^* = 0.65, *t*=7.77, *p*<0.0001

Quadratic:

(excluding potential contaminant) LMquadratic: *F*(2,36)=85.76, *R^2^_adj_*=0.82, *p*<0.0001

(including potential contaminant) LMquadratic: *F*(2,36)=84.85, *R^2^_adj_*=0.82, *p*<0.0001

Threshold Segmented:

(excluding potential contaminant)

A threshold model identified a changepoint in the slope at 33.2 (95% CI=28.1 to 57.0) copies (Fig. 1B): crosses with lower copy number have high heritability (*h_G_^2^*=1.60, *p*=<0.0001), whereas heritability was much lower (*h_G_^2^*=0.22, *p*<0.0001)

(including potential contaminant)

A threshold model identified a changepoint in the slope at 33.2 (95% CI=28.1 to 57.0) copies (Fig. 1B): crosses with lower copy number have high heritability (*h_G_^2^*=1.59, *p*=<0.0001), whereas heritability was much lower (*h_G_^2^*=0.22, *p*<0.0001)

Supplement C.

Incorporation of intra-individual variation in copy number to parental estimates of copy number, and estimates of number/proportion of offspring greater than this upward-adjusted parental sum.

| Cross_ID | Maternal_CN | Paternal_CN | mean_parental_CN | Max_intra_range | Max_intra_range_div2 |
| --- | --- | --- | --- | --- | --- |
| IG2 | 14.5 | 14.7 | 14.6 | 13 | 6.5 |
| C5 | 17.3 | 17.7 | 17.5 | 14 | 7 |
| IG6 | 19 | 16.7 | 17.85 | 16 | 8 |
| C9 | 25.2 | 20.2 | 22.7 | 13 | 6.5 |
| C6 | 24.9 | 30.8 | 27.85 | 13.5 | 6.75 |
| A9 | 22.6 | 33.6 | 28.1 | 14.5 | 7.25 |
| A3 | 43.1 | 16.9 | 30 | 17.5 | 8.75 |
| C7 | 38.2 | 23.6 | 30.9 | 14 | 7 |
| C2 | 38.3 | 23.8 | 31.05 | 14.5 | 7.25 |
| B4 | 32.5 | 30.8 | 31.65 | 19 | 9.5 |
| B10 | 36.5 | 29.9 | 33.2 | 16.5 | 8.25 |

…

| Mat_CN_intra_adj | Pat_CN_intra_adj | Parent_sumCN_intra_adj | Number_offspring_>_adjsum | Ppn_offspring_>_adjsum |
| --- | --- | --- | --- | --- |
| 21 | 21.2 | 42.2 | 0 | 0 |
| 24.3 | 24.7 | 49 | 3 | 0.15 |
| 27 | 24.7 | 51.7 | 2 | 0.0625 |
| 31.7 | 26.7 | 58.4 | 2 | 0.074074074 |
| 31.65 | 37.55 | 69.2 | 1 | 0.037037037 |
| 29.85 | 40.85 | 70.7 | 5 | 0.185185185 |
| 51.85 | 25.65 | 77.5 | 4 | 0.137931034 |
| 45.2 | 30.6 | 75.8 | 1 | 0.043478261 |
| 45.55 | 31.05 | 76.6 | 1 | 0.04 |
| 42 | 40.3 | 82.3 | 3 | 0.125 |
| 44.75 | 38.15 | 82.9 | 3 | 0.09375 |

Columns / variables:

Cross_ID = corresponds to first xx crosses in file ‘heritability_summary.csv’
Maternal_CN = estimated EPSPS copy number of maternal plant
Paternal_CN = estimated EPSPS copy number of paternal plant
Mean_parental_CN = mean of maternal and paternal copy number (parental midpoint copy number)
Max_intra_range = Average of the maximum intra-individual range between temporal replicates for the maternal and paternal plants (averaged across 2-5 intra-individual plants (intra_indiv_summary.csv) closest to parental CN)
Max_intra_range_div2 = previous column divided by two (here interested in the upper half)
Mat_CN_intra_adj = Maternal copy number plus intra-individual upward adjustment (Max_intra_range_div2)
Pat_CN_intra_adj = Paternal copy number plus intra-individual upward adjustment (Max_intra_range_div2)
Parent_sumCN_intra_adj = Sum of upward-adjusted maternal and paternal copy number
Number_offspring_>_adjsum = number of offspring with copy number greater than the upward-adjusted parental sum copy number
Ppn_offspring_>_adjsum = proportion of offspring with copy number greater than the upward-adjusted parental sum copy number
